# Transcriptional landscape of CD4+ T cells in Systemic Sclerosis

**DOI:** 10.64898/2026.01.03.697349

**Authors:** Gonzalo Villanueva-Martin, Gonzalo Borrego-Yaniz, Marialbert Acosta-Herrera, José Luis Callejas-Rubio, Norberto Ortego, Norbert Mages, Stefan Börno, Maria Gutierrez-Arcelus, Javier Martin, Lara Bossini-Castillo

## Abstract

**Objective:** This resource deeply characterizes the CD4+ T cell subpopulations populations in SSc patients, with a novel single-cell RNA sequencing (scRNA-seq) approach, offering unprecedented insights into the cellular and molecular underpinnings of the disease.

**Methods:** We performed scRNA-seq to analyze over 80,000 CD4+ T cells from 8 SSc patients and 8 healthy controls, integrating abundance, transcriptional, and TCR repertoire analysis, to define CD4+ T cell subtype-specific signatures associated with SSc.

**Results:** SSc CD4+ T cells displayed a global interferon-driven activation signature and significantly reduced TNFα signaling. Key transcriptomic alterations included the downregulation of *SOX4* and *CD83*, alongside an expansion of Th2 and proinflammatory, steroid-resistant Th17 cells. Furthermore, Tregs exhibited a destabilized state characterized by high *FCRL3* and reduced *FOXP3* expression. Finally, TCR repertoire analysis identified increased clonal expansion specifically within the central memory (Tcm) compartment.

**Conclusion:** Together, this data revealed SSc-associated cell population states in CD4+ T cells with unique transcriptomic signatures and provided a publicly accessible interactive platform to facilitate exploration of this dataset by the research community.

## Introduction

Systemic sclerosis or scleroderma (SSc), is a chronic immune-mediated inflammatory disease (IMID). The pathogenesis of SSc involves an intricate interplay of immune cell dysfunction, vascular damage, and progressive skin fibrosis ^1^. Additionally, SSc predominantly affects middle-aged women ^1^.

Clinical and genetic evidence underscores a significant disruption of the immune system in the development of SSc, with a particularly notable involvement of lymphocyte activation and T cell subsets ^2,3^. Among these, CD4+ T cells play a pivotal role not only as regulators of the immune response but also as effectors contributing to inflammation and tissue damage ^2,3^. Additionally, the association of more than 30 genetic loci with SSc ^4^, many of which are related to T cell activation, highlights the importance of studying the CD4+ T cell lineage to better understand SSc susceptibility and to develop more targeted therapies.

Systemic immune deregulation is a SSc hallmark, but no increase in absolute cell numbers has been observed to date in the different subsets of circulating CD4+ helper T cells (Th) ^5^. However, an imbalance in Th1 and Th2 cytokine production is well-established in SSc patients ^5^. Furthermore, a skewed differentiation of Th cells towards the proinflammatory Th17 phenotype, as opposed to regulatory T cells (Tregs) that control the immune response, has been recognized as a feature of SSc ^6,7^. Although, Th17 cells have been implicated in the pathogenesis of SSc, particularly in vascular damage ^5^, reports on Th17 abundance in SSc peripheral blood (PB) are inconsistent ^8^. Despite this controversy, imbalanced Th17 and Treg activity is considered critical in SSc pathogenesis. Altered proportions of Tregs in the PB and affected tissues of SSc patients and aberrant SSc-related gene expression have been reported, further supporting the relevance of this imbalance ^9^.

Additional scarce T cell subsets have also shown characteristic alterations in SSc ^5^. For instance, the Th22 subset, known for its profibrotic role in tissue repair, wound healing, and autoimmunity, was overrepresented in SSc-affected skin tissues ^10^. Moreover, a SSc-specific recirculating T helper follicular cell (Thf) subset, defined by high *CXCL13* expression (CXCL13+Thf), was identified in the skin tissue of SSc patients ^11^ suggesting that the study of scarce CD4+ T cell subsets in SSc might shed light on the etiopathogenesis of this IMID.

A key element in T cell biology is the T cell receptor (TCR), which is a highly polymorphic and antigen-specific membrane receptor required for T cell activation ^12^. Lower diversity in the TCR repertoire of IMID patients is frequently observed ^13^, and it was recently described in SSc patients in comparison to healthy donors ^14^. However, little is known about the TCR repertoire in specific CD4+ T cell subtypes and its association with SSc-related aberrant activation and behavior.

Considering the accumulating evidence supporting a central role of circulating CD4+ T cells in SSc pathogenesis, we performed a comprehensive single-cell characterization of the transcriptomes of peripheral blood (PB) CD4+ T cells from patients with SSc and unaffected controls. We established a publicly accessible transcriptomic resource comprising more than 80,000 CD4+ T cells and their TCR repertoires, enabling detailed interrogation of cellular heterogeneity and clonal architecture within this compartment.

## Materials and methods

### Patient description

This study included a cohort of 16 female participants, all with self-reported European ancestry. The average age was 60 years old for SSc patients and 59 for the unaffected controls. Among the participants, 8 were diagnosed with SSc according to the diagnostic criteria proposed by the ACR ^15^, and they were further classified, according to the criteria proposed by LeRoy, into limited cutaneous (lcSSc) or diffuse cutaneous SSc (dSSc) ^16,17^. Participants were recruited at Hospital Universitario San Cecilio, Granada, Spain, with informed consent. Clinical information, including anonymized serological profiles and drug treatment of the patients, is shown in **Supplementary Table 1**.

### Cell suspension protocol

PB mononuclear cells (PBMCs) were separated as previously described ^18^. Using a magnetic bead kit (Stem Cell Easy Step, ref #17858) CD4+ cells were negatively selected, following manufacturer’s protocol. Flow cytometry and mRNA expression confirmed CD4+ cell purity (>99.6%, **Supplementary Figure 1**). Subsequently, CD4+ cell suspensions were cryopreserved as described elsewhere ^18^.

### Single cell RNA-sequencing and T cell receptor library generation

For single-cell whole transcriptome sequencing (scRNA-seq), the Next GEM technology provided by 10x Genomics was selected. Following the manufacturer’s instructions, the samples were processed using the Chromium Next GEM Chip G Single Cell Kit (10xGenomics, PN-1000127 for gene expression and PN-1000005 for V(D)J profiling) and Chromium Next GEM Single Cell 5’ Library and Gel Bead Kit v1.1 (10xGenomics, PN-1000165_a). Subsequently, the generated cDNA libraries were sequenced on the NovaSeq 6000 platform (Illumina) using S2 and SP chemistry v1.5. For the gene expression, these settings allowed an average 74.32% of reads assigned to cells, with an average of 4,372 UMIs per cell. For the TCR immune profiling, an average of 70.71% of reads in cells was observed. The sequencing reads were aligned to the GRCh38 genome build, and the unique molecular identifiers (UMI) were processed using the 10x Cell Ranger Single Cell Software Suite (v3.0.0) with default parameters, resulting in an average of 1,357 genes identified per cell.

### Single cell RNA-sequencing data analysis

The outcomes from Cell Ranger were imported into *Scanpy* (v1.8.2) ^19^ within a Python environment (v3.8.1). All samples successfully met the established quality control (QC) filters: cells with fewer than 500 genes detected, with over 10% of reads mapped to mitochondrial genes, or having more than 30% of reads aligned to ribosomal genes were systematically excluded. Furthermore, to preclude potential doublets, cells with more than 3,000 detected genes were removed. Finally, 81,714 CD4+ T cells and 21,583 genes passed all the QC filters.

A Batch Balanced k-Nearest Neighbour (BBkNN) integration graph was calculated using the first 20 principal components and clustering was performed employing the Leiden algorithm. No batch effect between SSc patients and controls were observed either in the PCA or UMAP embeddings. The validity of the integration was supported by consistent clustering outcomes observed through alternative algorithms and the analysis of the UMAP representations of each individual after integration (**Supplementary Figure 2A**).

Afterwards, unsupervised clustering by the Leiden algorithm returned 12 clusters, which were assigned to known CD4+ cell subtypes based on recent markers described in scRNA-seq studies in CD4+ cells ^20^ (**Figure 2A**). However, each cluster was analyzed as a distinct entity, independently of the previously assigned cell subtype.

We performed differential gene expression using a pseudobulk approach with a lineal model for the global analysis (all CD4 T+ together, patients vs. controls) and the Wilcoxon rank-sum statistical test with Scanpy for the clusters and annotation and per cluster differential expression between patients and controls. Additionally, Gene Set Enrichment Analysis (GSEA) was performed to determine the pathway enrichment with the fgsea function from fgsea R package (v1.32.2)^21^. Transcriptomic Factor (TF) Activity was calculated using the *run_ulm* function from decoupleR (v2.12.0)^22^. Statistical significance was defined after applying the Benjamini-Hochberg False Discovery Rate correction method. We applied Partition-based Graph Abstraction to test the statistical strength of the connectivity between the clusters. To identify cell differentiation trajectories, we used the *Monocle3* algorithm ^23^.

Finally, *Scirpy* (v1.0) was used for the analysis of TCR clonotypes and clonotype networks, clonal expansion, normalized Shannon entropy, TCR repertoire, preferential usage of V(D)J genes and epitope analysis.

Details of the scRNA-seq analyses are found in the **Additional Information** file

## Results

### CD4+ T cells from SSc patients exhibit IFN-mediated activation profiles

CD4+ T lymphocytes were extracted from the PB of 8 women with SSc diagnosis, alongside 8 unaffected women (CTRL) matched by age and ethnicity (**Figure 1A**), as detailed in **Supplementary Table 1**.

**Figure 1.**
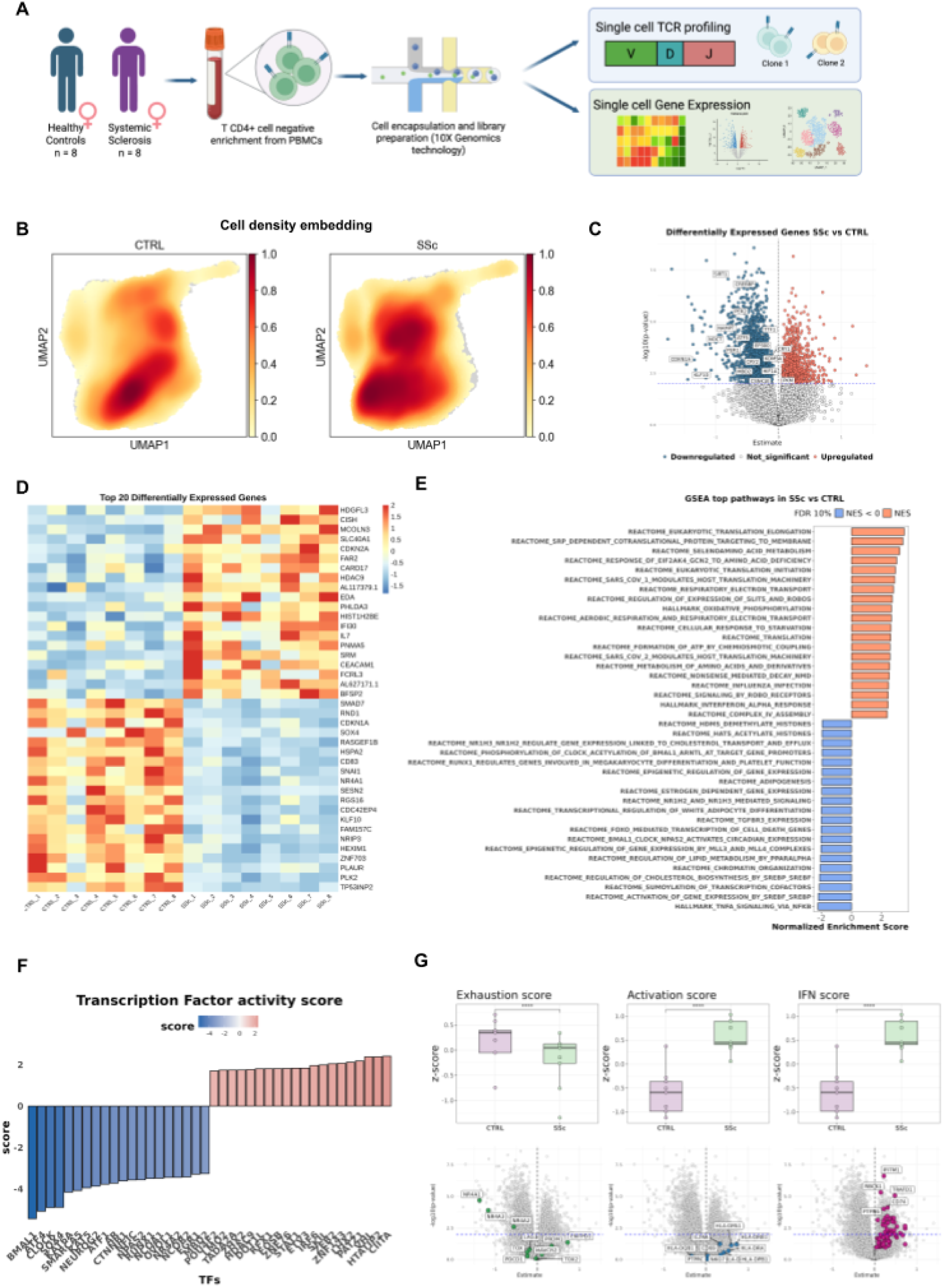
Single-cell characterization of circulating CD4+ T cells in systemic sclerosis (SSc) and controls (CTRL). **(A)** Study design. PB CD4+ T cells were isolated from 8 SSc patients and 8 age-matched female CTRL and profiled using 10X Genomics chemistry, enabling simultaneous gene expression (GEX) and TCR repertoire analysis. All processed data are available through an interactive Shiny application. **(B)** UMAP visualization showing cell density distribution by condition (CTRL and SSc). **(C)** Volcano plot of differentially expressed genes (DEGs) identified by global pseudobulk analysis. Upregulated genes (FDR < 0.05; L2FC > 0) are shown in red, downregulated genes (FDR < 0.05; L2FC < 0) in blue, and non-significant genes in grey. The top 20 upregulated and downregulated genes are labeled. **(D)** Donor-level heatmap of the top 20 upregulated and top 20 downregulated genes identified in the pseudobulk analysis. **(E)** Gene Set Enrichment Analysis (GSEA) using Reactome and Hallmark databases based on pseudobulk results. Pathways with normalized enrichment score (NES) > 0 are shown in orange and NES < 0 in blue. **(F)** Transcription factor (TF) activity scores inferred using decoupleR. TFs with positive activity scores are shown in orange and negative scores in blue. **(G)** Exhaustion, activation, and IFN scores derived from predefined gene signatures. Scores represent the average expression of genes within each signature. Boxplots show per-condition distribution, and volcano plots display the direction and significance of individual genes contributing to each score.

Following QC, a final set of 81,637 CD4+ high quality scRNA-seq transcriptomes of individual cells from both SSc and unaffected individuals were selected for analysis. Although the number of cells was balanced between the patients with SSc and in the CTRL (∼4,100 per SSc patients and ∼4,000 for CTRL, **Supplementary Table 2**), the cell density on a UMAP embedding was not uniform across patients and controls and had clearly distinct patterns (**Figure 1B**). We hypothesized that these differences in cell density reflect variations in T cell states between SSc patients and controls.

The pseudobulk differential expression analysis (DEA) between SSc and control samples (**Supplementary Table 3**) identified 2,371 differentially expressed genes (DEGs) at FDR < 0.05, including 895 upregulated and 1,476 downregulated genes in SSc (**Figure 1C**). Among the top upregulated DEGs, we observed multiple IFN-responsive genes, including *IFITM1*, *IFITM2*, *IFI30*, and *CISH*, as well as *FCRL3*, which has been implicated in Treg and Th17 differentiation (**Figure 1C and 1D; Supplementary Table 3**). Conversely, among the most downregulated genes were *SOX4*, a negative regulator of T helper 2 (Th2) differentiation^24^ but positive for T follicular helper (Thf) development^25^, and *CD83*, a regulator of T regulatory T cells (Treg) development (**Figure 1C and 1D**). Notably, *NR4A1* and *NR4A2*, members of the Nuclear Receptor Subfamily 4 Group A, were markedly downregulated in SSc T CD4+ cells (**Supplementary Table 3**). These genes are typically associated with T cell exhaustion when expressed at high levels^26,27^.

To characterize the biological processes associated with these transcriptional changes, we performed Gene Set Enrichment Analysis (GSEA) using the Reactome and Hallmark databases (**Figure 1E**), together with Transcription Factor (TF) activity inference using decoupleR^22^ (v2.9.7) (**Figure 1F**). GSEA revealed significant enrichment of IFN-α and IFN-γ signaling pathways in SSc samples, along with pathways related to mitochondrial energy metabolism. In contrast, pathways associated with lipid metabolism and chromatin organization were negatively enriched (**Figure 1E**), along with a negative enrichment of TFs involved in the circadian rhythm and Treg development, such as *RUNX1* (**Figure 1F**).

Considering the pathway enrichment and TF activity results, we further quantified transcriptomic signatures related to exhaustion, activation, and IFN responses. As shown in **Figure 1G**, SSc T CD4+ cells displayed increased IFN and activation scores in patients compared to controls, while exhaustion scores were not elevated (**See Additional information**). Importantly, although the activation signature was globally increased as an expression profile, none of the included genes reached the predefined FDR threshold in the DEA individually. Therefore, our results suggest that the SSc T CD4+ cells have an IFN-driven activated but not exhausted profile in comparison with the healthy controls.

### Altered activation signatures characterize SSc CD4+ clusters

We identified 12 distinct CD4+ T cell clusters, none of which were exclusively composed of either SSc or CTRL cells (**Figure 2A**). Among these, three clusters displayed differential abundance between patients and controls: cluster 3 showed an increased proportion of SSc cells, whereas clusters 4 and 9 were reduced in SSc compared to CTRL (**Figure 2B**).

**Figure 2.**
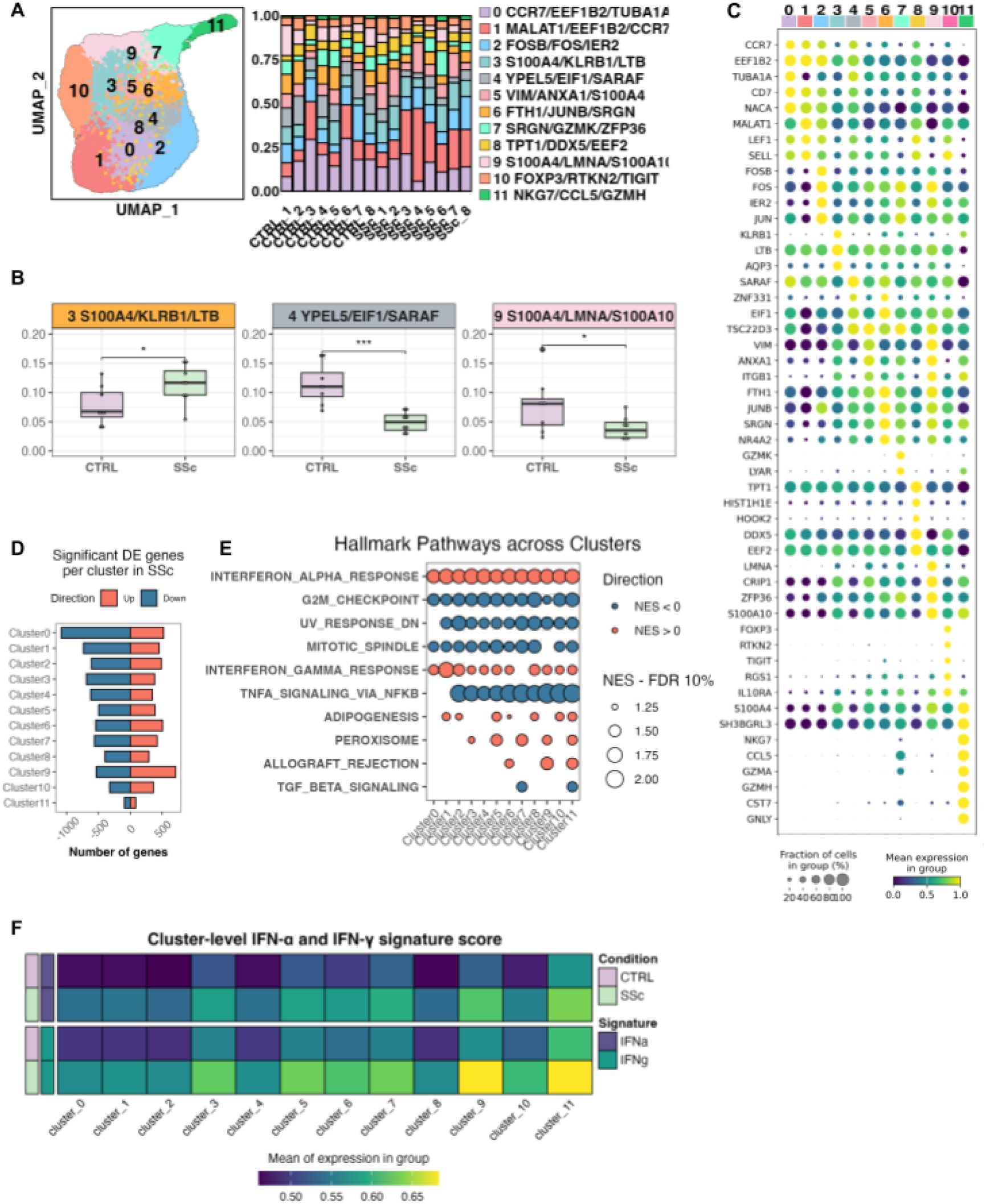
Clustering and transcriptional characterization of CD4+ T cell populations in SSc. **(A)** UMAP visualization of CD4+ T cell clusters and compositional analysis. Twelve clusters (0–11) were identified and labeled according to relative size. The stacked bar plot shows the contribution of each individual donor to each cluster. The top three marker genes defining each cluster are indicated. **(B)** Boxplots showing differences in cluster abundance between SSc patients and CTRLs (SSc in light green; CTRL in light purple). Significance was assessed using a Wilcoxon rank-sum test with FDR correction. *FDR ≤ 0.05; **FDR ≤ 0.001. **(C)** Dot plot of cluster-defining marker genes. Dot size represents the percentage of cells expressing the gene, and color intensity indicates mean expression level within each cluster. **(D)** Number of significantly differentially expressed genes (DEGs) per cluster identified by Wilcoxon rank-sum test comparing SSc and CTRL cells. Upregulated genes (FDR < 0.05 and Z-score > 0) are shown in orange and downregulated genes (FDR < 0.05 and Z-score < 0) in blue. **(E)** Hallmark pathway enrichment across clusters based on pseudobulk DEGs and cluster-level Wilcoxon analysis. Pathways with normalized enrichment score (NES) > 0 are shown in orange and NES < 0 in blue. Dot size reflects significance at FDR < 10%. **(F)** Cluster-level heatmap of IFN-α and IFN-γ signature scores by condition (SSc in light green; CTRL in light purple). Color intensity represents the averaged leading-edge gene expression score per cluster.

We applied the Wilcoxon rank-sum test to define the transcriptomic signature of each cluster (**Figure 2C; Supplementary Table 4**) and to assess differential expression between SSc and CTRL cells within each cluster (**Figure 2D**). Across clusters, SSc cells showed more downregulated genes and fewer upregulated genes relative to CTRL cells, with the exception of cluster 9 (**Figure 2D; Supplementary Table 5**).

To further refine the IFN signal identified in the pseudobulk analysis, we performed GSEA on the cluster-level differential gene expression results. With the exception of IFN-γ signaling in cluster 7 all clusters showed enrichment in IFN-α and IFN-γ signalling pathways (**Figure 2E**). Furthermore, we calculated the score for each cluster in each condition, by computing the average expression of the leading edge genes for both pathways per cell and subsequently averaged across cells within each cluster and condition, showing higher scoring in SSc patients’ cells in every cluster, especially in clusters 9 and 11 (**Figure 2F**). In contrast, pathways related to cell proliferation, including G2M checkpoint and mitotic spindle signatures, were negatively enriched across clusters, together with TNFα signaling via NFKB. NFKB is a central regulator of T cell activation and differentiation, and its dysregulation has been implicated in autoimmunity^28^. The coordinated reduction of TNFα–NFKB signaling across clusters suggested a broad alteration of activation-associated transcriptional programs in SSc CD4+ T cells.

### Annotation of CD4+ T cell clusters reveals canonical differentiation states

Considering the different cluster marker genes (**Figure 2C** and **Supplementary Table 4**), each cluster was classified according to their responsiveness to activation, proliferative capacity and effector potential ^20^ into: naive T cells (Tn), central memory T cells (Tcm), effector memory T cells (Tem), re-expressing *CD45RA* effector memory T cells (Temra) (also known as CD4+ cytotoxic T Lymphocytes, CD4+ CTL), and Treg (**Figure 3A**).

**Figure 3.**
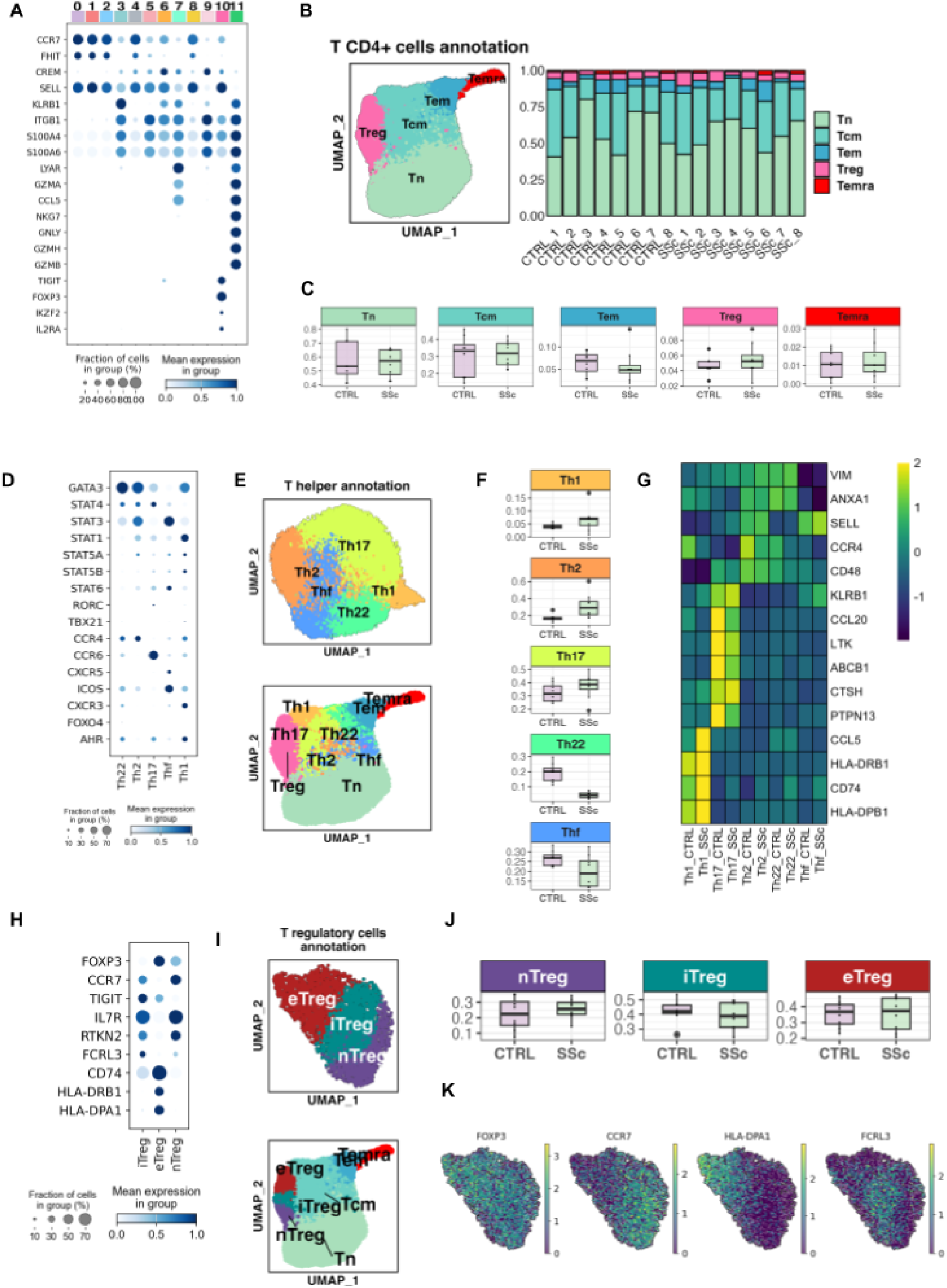
Annotation of CD4+ T cell subsets and subclustering of Tcm and Treg compartments. **(A)** Dot plot of canonical marker genes used to annotate the 12 CD4+ T cell clusters based on established lineage-defining markers. Dot size represents the fraction of expressing cells and color intensity indicates mean expression. **(B)** UMAP visualization and compositional analysis after cell type annotation. CD4+ T cells were classified as Tn (naive T cells), Tcm (central memory T cells), Tem (effector memory T cells), Temra (CD45RA-re-expressing effector memory T cells), and Tregs (regulatory T cells). The stacked bar plot shows the relative contribution of each individual to each annotated subset. **(C)** Boxplots showing the relative abundance of annotated CD4+ T cell subsets in CTRL and SSc samples. **(D)** Dot plot of marker genes used to define T helper (Th) subpopulations within the Tcm compartment. **(E)** UMAP visualization of Tcm subclustering and subsequent annotation into Th1, Th2, Th17, Th22, and Thf subsets. The lower panel shows projection of Th annotation onto the global CD4+ T cell UMAP. **(F)** Boxplots showing the distribution of CTRL and SSc cells across Th subclusters. **(G)** Heatmap of key differentially expressed genes across Th subsets, highlighting lineage-specific and disease-associated transcriptional programs. **(H)** Dot plot of canonical marker genes used to define Treg subpopulations. **(I)** UMAP visualization of Treg subclustering and projection onto the global CD4+ T cell UMAP. **(J)** Boxplots showing the relative abundance of nTreg, iTreg, and eTreg subsets in CTRL and SSc samples. **(K)** UMAP feature plots of FOXP3, CCR7, HLA-DPA1, and FCRL3 expression across Treg subclusters.

As a result of this classification (**Figure 3B**), 5 clusters were identified as Tn, encompassing 57.3% of the total cells. These clusters exhibited the highest expression of *CCR7*, a hallmark Tn marker. By the combined expression of *CREM*, *ITGB1*, and *KLRB1* among others, 4 clusters (30.2% of the cells) were classified as Tcm. Cluster 7 (6.1% of the cells) was identified as Tem, due to a high expression of *CCL5* and *GZMA*. Cluster 10 (5.1% of all cells) showed a high *FOXP3* expression, and it was identified as Tregs. Lastly, cluster 11 (1.2% of cells) was classified as Temra/CD4+ CTL, with elevated expression of *GZMH* and *GZMB*. We observed no statistically significant differences in the abundance of the different T cell subtypes neither between patients and healthy controls (**Figure 3C**).

Trajectory and PAGA analyses supported the cell type annotation (**Supplementary Figure 2C**). PAGA showed strong connectivity among Tn and Tcm clusters, while single-cluster cell types appeared more isolated. Diffusion mapping identified two main branches originating from cluster 0: one leading to highly differentiated Temra cells (cluster 11) with the highest pseudotime, and another comprising a combination of Tcm and Tem cells.

### CD4+ T cell subtypes exhibit altered immune activity in SSc

Giving the heterogeneity of the Tcm compartment and its relevance for IMIDs ^29^, we sought to further characterize the composition of CD4+ T cell subtypes. To accomplish this, the clusters annotated as Tcm (clusters 3, 5, 6, and 9; comprising a total of 24,708 cells) were selected and re-clustered. This analysis identified nine sub-clusters with a homogenous distribution between SSc and Temra/CTL. Using a panel of established T helper cells (Th) markers, we classified these clusters as: Th1 marked by high expression of *STAT1, CXCR3* and *AHR*; Th2 marked by expression of *GATA3*, *STAT3* and *STAT4*; Th17 marked by expression of *RORC* and *CCR6*; Th22 expressing *GATA3* and *ICOS*; and Thf expressing *CXCR5* and *ICOS* (**Figure 3D-E; Supplementary Table 6**).

The analysis of the cellular composition revealed significant differences between patients and controls. SSc samples showed an increased abundance in Th2 cells and a reduction in the Th22 subset (**Figure 3F**). Th2 cells exhibited the expression of *XBP1* ^30^, as activation markers like *SELL*, *CCR4*, and *CD48*/*SLAMF2* (**Figure 3G**; **Supplementary Table 6**). Notably, *SOX4* was downregulated in SSc Tn cells, consistent with the global pseudobulk analysis. *SOX4* has been reported to negatively regulate Th2 differentiation through direct interaction with GATA3^24^. Therefore, reduced *SOX4* expression in Tn cells may facilitate Th2 polarization and could contribute to the increased abundance of this subset observed in SSc. These findings support a highly activated SSc-specific transcriptional profile in circulating Th2 cells.

The Th17 population was predominantly contained within cluster 3, with 85.6% of the Th17 annotated cells coming from that cluster, which was expanded in SSc (**Figure 3E**; **Supplementary Table 4** and **Figure 3F**). Th17 cells displayed a pronounced proinflammatory signature characterized by the expression of *KLRB1/CD161* and *CCL20* (**Figure 3G**; **Supplementary Table 6**). In addition, previously reported markers of pathogenic glucocorticoid-resistant Th17 activation were also increased in this subset, for example, *LTK*, *ABCB1* (also known as MDR1 or P-glycoprotein), *CTSH*, and *PTPN13* (**Figure 3G; Supplementary Table 6**).

Th1 and Th22 populations were mainly represented within cluster 9, which was found to be underrepresented in SSc patients (**Figure 2C**). Notably, cells from SSc patients exhibited a biased towards effector memory characteristics, characterized by the increased expression of *CCL5* and MHC class II genes, *i.e*. *HLA-DRB1* and *HLA-DPB1* (**Figure 3G; Supplementary Table 5)**. SSc cells within these clusters, as well as Temra cells exhibited high IFN-α and IFN-γ scores in contrast with other clusters or cell annotations, very similar as Temra cells showed (**Figure 2F**). Finally, the elevated expression of the MHC class II genes in this cluster goes in line with the increased inferred CIITA TF activity observed in SSc, which was primarily driven by these cells (**Figure 1F**).

### Tregs population express markers of dysregulation in SSc

Cluster 10 was identified as Treg cells based on canonical markers. Within this population, *TIGIT* expression was markedly increased (**Figure 2B**), consistent with previous reports in SSc, describing elevated levels of this gene in patients Tregs ^31^. In parallel, *FCRL3* was significantly upregulated in SSc both at the pseudobulk levels (**Figure 1C–D**) and within the Treg resolution (**Supplementary Table 5**). *FCRL3* encodes Fc Receptor Like 3, an immunoglobulin receptor whose stimulation has been associated with attenuation of Treg suppressive function ^32^, suggesting that its increased expression may have functional consequences in the SSc context.

To examine Treg heterogeneity, we performed subclustering of cluster 10. Using established Treg markers (**Figure 3H**), we identified three subpopulations: naive Treg (nTreg), characterized by high expression of *CCR7* and *IL7R*; intermediate Treg (iTreg), expressing intermediate levels of *CCR7* and *CD74* and elevated *TIGIT*; and effector Treg (eTreg), defined by low *CCR7* and increased expression of *HLA-DRB1*, *HLA-DPA1*, and *CD74* (**Figure 3H–I**). No significant differences in subtype abundance were observed between SSc and CTRLs (**Figure 3H-I**), indicating that transcriptional remodeling rather than compositional shifts characterizes the Treg compartment in SSc.

Notably, the iTreg subset displayed the highest *FCRL3* expression of all cell subsets together with reduced *FOXP3* levels (**Figure 3K**). This combination of elevated inhibitory receptor expression and diminished lineage-defining transcription factor levels suggests a destabilized regulatory state.

### Tcm cell clones are differentially expanded in SSc CD4+ T cells

To assess T-cell receptor usage, an immunological profiling of cells was conducted for both patients and controls. This profile aimed to examine differences in terms of clonal expansion (an increment in the abundance of a specific clonotype) and V(D)J gene usage.

There was no observable global difference in T cell expansion between SSc and CTRL in the clonotype expansion (**Figure 4A**). Considering the cell subtypes, the most differentiated subtypes i.e. Tcm, Tem and Temra subtypes showed an increased expansion (**Figure 4A**), but differences between SSc and controls were only significant in the Tcm subtype. The clonal expansion analysis by cluster revealed significant increases in clusters 3, 5, 7, 8, 9, and especially in cluster 11 (FDR < 0.05) (**Figure 4B**). However, differences in clonal expansion between SSc and controls only appeared in clusters 3 and 5 (FDR < 0.05) compared to other clusters. Nevertheless, Shannon diversity measures did not significantly differ between SSc and CTRL in any of the CD4+ T cell clusters or subtypes. The V(D)J analysis showed no bias in the usage of these genes between SSc patients and CTRL (**Figure 4C**), nor between SSc subtypes. As for the epitope analysis, of 81,637 cells, only ∼9,000 cells were matching successfully with an epitope in VDJdb ^33^. More than 20 different epitopes were inferred for all the cells, but no relevant differences in the epitope recognition were observed between SSc patients and the non-affected CTRL (**Figure 4D**).

**Figure 4.**
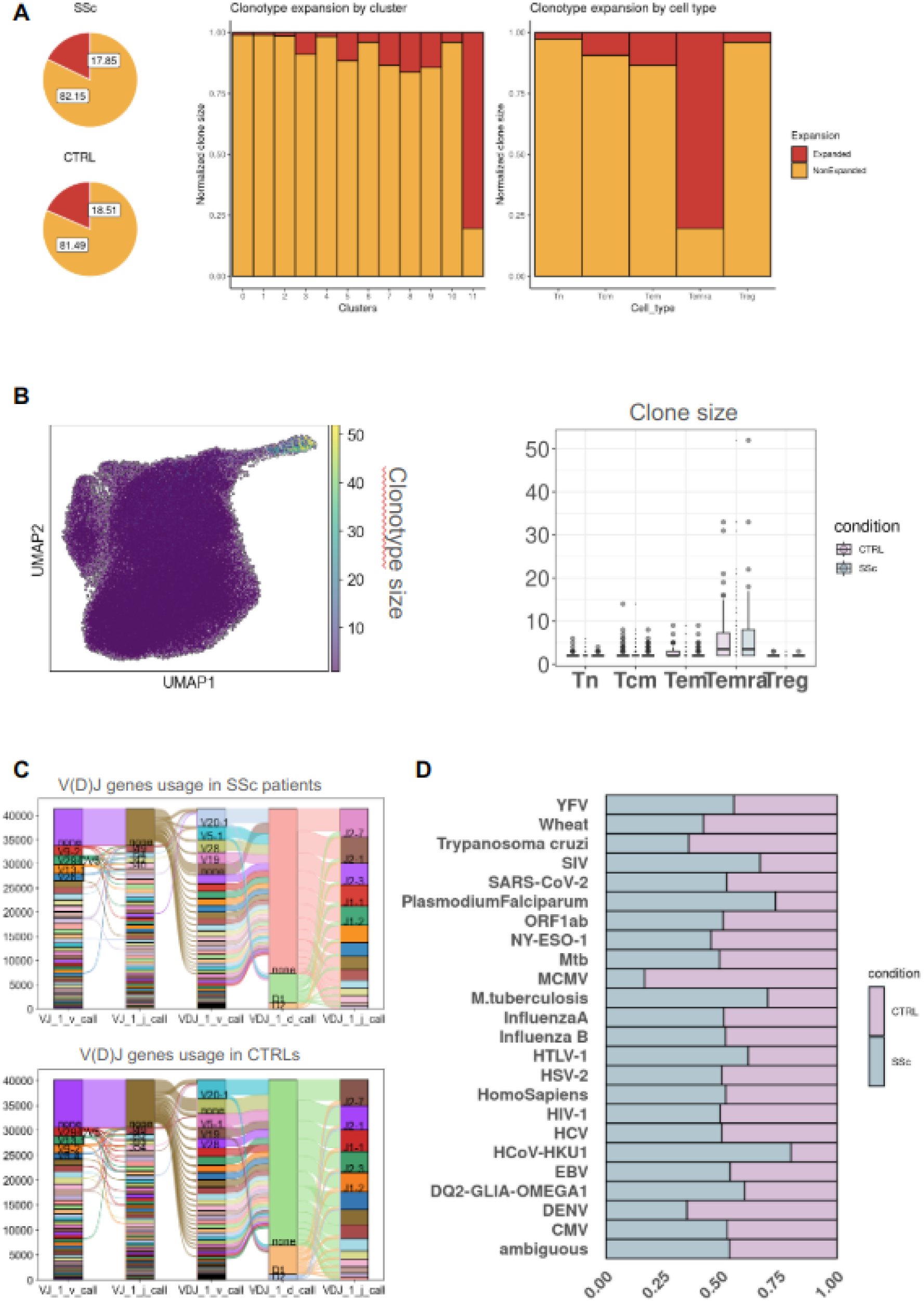
Clonotype expansion and TCR repertoire analysis in CD4+ T cells. **(A)** Clonotype expansion analysis. Left: Visualization of clonotype expansion across all cells for SSc and CTRL. Middle: Comparison of clonotype expansion between SSc and CTRL by cluster. Right: Comparison of clonotype expansion between SSc and CTRL by cell type. **(B)** UMAP visualization and clonotype size distribution. Left: UMAP plot colored by clonotype size, showing the distribution of expanded clonotypes across the CD4+ T cell population. Right: Box plot displaying the distribution of clonotype sizes across different cell types. **(C)** VDJ gene usage analysis. Ribbon plots illustrating the usage of VDJ genes in the TCR repertoire: Top panel: VDJ gene usage in SSc patients. Bottom panel: VDJ gene usage in CTRL. **(D)** Epitope analysis. Visualization of epitopes identified within the single-cell subset, highlighting potential antigen-specific responses in SSc and CTRL.

## Discussion

Here, we provide a single-cell transcriptomic resource of circulating CD4+ T cells in peripheral blood (PB) from patients with SSc and unaffected controls. By integrating compositional, transcriptional, and TCR analyses, this dataset defines subtype-specific transcriptional states and reveals disease-associated remodeling across regulatory and effector CD4+ compartments. While most previous scRNA-seq studies in SSc have focused on affected tissues^11,34–36^, our work establishes a complementary PB-based framework to investigate systemic immune alterations. This resource enables direct comparison between circulating and tissue-resident CD4+ T cell states and supports future integrative analyses of immune dysregulation in SSc.

The proportion of Tregs in PB from SSc patients remains controversial, with reports describing either increased frequencies^37,38^ or reduced numbers in skin but not blood^39^. In our dataset, Treg abundance did not differ between SSc and CTRL (**Figure 3B-J**); however, transcriptional remodeling was noticeable. SSc Tregs displayed elevated *FCRL3* expression, particularly within the intermediate Treg (iTreg) subset (**Figure 3H**). *FCRL3* signaling has been associated with reduced suppressive capacity and acquisition of Th17-like features^32^. Subclustering identified naive, intermediate, and effector Treg states without compositional differences between patients and controls, suggesting that transcriptional changes rather than abundance shifts characterize the Treg compartment in SSc. The iTreg subset exhibited high *FCRL3* and reduced *FOXP3* expression, consistent with altered lineage stability. Although *TIGIT* upregulation has been linked to T cell exhaustion in SSc^31^, our data do not support a generalized exhaustion phenotype (**Figure 1G**). *TIGIT* was elevated within the Treg population but not across the broader CD4+ cells. Moreover, despite the reported link between *TIGIT* signaling and *AREG*-mediated tissue repair^40,41^, *AREG* expression was not increased in SSc Tregs (**Supplementary Table 4**). Given the known plasticity between Treg and Th17 programs^42^, this transcriptional configuration may reflect a partial switch and reduced suppressive capacity. Together with the global IFN-driven signature and transcription factor instability in SSc, these findings support the hypothesis in which Treg are transcriptionally and functionally compromised rather than classical exhaustion, contributing to the general immune dysregulation in patients.

We also identified expansion of a Th17-containing cluster in SSc. These cells expressed proinflammatory markers including *KLRB1*/*CD161* and *CCL20* (**FIgure 2C and 3G**), consistent with a migratory and inflammatory phenotype described in other immune-mediated diseases^43–45^. This observation suggests that circulating Th17 subsets may contribute to systemic immune activation in SSc.

TCR analysis showed increased clonal expansion in Tcm from SSc, but no other significant differences. The lack of clear differences between SSc patients and controls may be attributed to the strong heterogeneity of the patients, the sample size, or the limitations of the tools used. Nevertheless, further analyses are required to confirm the role of TCR diversity in SSc.

Our study has several limitations. DEA based on Wilcoxon rank-sum testing may be influenced by single-cell inter-dependence and patient heterogeneity, leading to inflation of low P values. Treatment exposure, and comorbidities may introduce additional variability. Although no individual or subgroup drove the observed patterns, larger cohorts will be necessary to refine subtype-specific signatures. TCR analysis revealed increased clonal expansion within Tcm cells in SSc but no broad differences in repertoire diversity, warranting further investigation.

In conclusion, this study provides a single-cell transcriptional resource of PB circulating CD4+ T cells in SSc, defining CD4+ T cell subtype-specific transcriptional states, identifying remodeling within the Treg compartment, and highlighting expansion of Th2 compartment and the proinflammatory Th17 populations. By making these data available, we aim to facilitate integrative analyses across tissues and diseases and to support future efforts to dissect immune dysregulation in SSc.

## Acknowledgements

We would like to thank Sofia Vargas and Gemma Robledo for their excellent technical assistance, and also to Sven Klages for his excellent bioinformatic assistance. We also appreciate the controls and the affected individuals who generously provided the samples for these studies.

## Contributors

GVM: data analysis, data interpretation, manuscript drafting, revision and approval; GBY: data analysis, manuscript revision and approval; MAH: study design, revision and approval; NOC: data acquisition, manuscript revision and approval; JLCR: data acquisition, manuscript revision and approval; NM: experiments and manuscript revision and approval; SB: data analysis, manuscript revision and approval; MGA: analysis design, manuscript drafting; JM: study design, data interpretation, manuscript drafting, revision and approval; LBC: study design, data interpretation, manuscript drafting, revision and approval.

## Ethical approval

An ethical protocol was prepared with consensus across all partners and was approved by the local ethical committee of the clinical recruitment center. The study adhered to the standards set by the International Conference on Harmonization and Good Clinical Practice (ICH-GCP), and to the ethical principles that have their origin in the Declaration of Helsinki (2013). The protection of the confidentiality of records that could identify the included subjects is ensured as defined by the EU Directive 2001/20/EC and the applicable national and international requirements relating to data protection in each participating country. This study was approved by the Ethical Committee “CEIM/CEI Provincial de Granada (0371/2020)”, and the CSIC Ethical Committee (026/2020).

## Data availability

The raw counts, barcode and matrix, normalized counts and metadata are available in GEO. Additionally, anonymized demographic information, differential expression results, pathway enrichment results, and data visualization is available in the publicly open ShinyApp, https://mgalab.shinyapps.io/CD4inSScResource/.

## Code availability

All the codes used for processing, analyzing and visualizing the data in this study were compiled into a single publicly available GitHub repository: https://github.com/gonv/SSc_CD4_project.

